# Histone acetyltransferase HAF2 associates with PDC to control H3K14ac and H3K23ac in ethylene response

**DOI:** 10.1101/2023.12.31.573642

**Authors:** Chia-Yang Chen, Zhengyao Shao, Guihua Wang, Bo Zhao, Haley A. Hardtke, Josh Leong, Tiffany Zhou, Y.Jessie Zhang, Hong Qiao

## Abstract

Ethylene plays its essential roles in plant development, growth, and defense responses by controlling the transcriptional reprogramming, in which EIN2-C-directed regulation of histone acetylation is the first key-step for chromatin to perceive ethylene signaling. However, the histone acetyltransferase in this process remains unknown. Here, we identified histone acetyltransferase HAF2, and mutations in HAF2 confer plants with ethylene insensitivity. Furthermore, we found that HAF2 interacts with EIN2-C in response to ethylene. Biochemical assays demonstrated that the bromodomain of HAF2 binds to H3K14ac and H3K23ac peptides with a distinct affinity for H3K14ac; the HAT domain possesses acetyltransferase catalytic activity for H3K14 and H3K23 acetylation, with a preference for H3K14. ChIP-seq results provide additional evidence supporting the role of HAF2 in regulating H3K14ac and H3K23ac levels in response to ethylene. Finally, our findings revealed that HAF2 co-functions with pyruvate dehydrogenase complex (PDC) to regulate H3K14ac and H3K23ac in response to ethylene in an EIN2 dependent manner. Overall, this research reveals that HAF2 as a histone acetyltransferase that forms a complex with EIN2-C and PDC, collectively governing histone acetylation of H3H14ac and H3K23ac, preferentially for H3K14 in response to ethylene.

## Introduction

Ethylene is a pivotal plant hormone that plays pleiotropic roles in plant growth, development, stress response and defense response to pathogens by controlling downstream gene transcription, protein translation and posttranslational modifications, chromatin remodeling, and epigenetic regulation of gene expression^1–12^. The ethylene signal is perceived by ethylene receptors residing on the endoplasmic reticulum (ER) membrane^13,14^. In the absence of ethylene, the ethylene receptors and CONSTITUTIVE TRIPLE RESPONSE 1 (CTR1), an ER membrane associated kinase that directly interacts with one of ethylene receptors ETHYLENE RESPONSE1 (ETR1), are activated^15,16^. CTR1 phosphorylates ETHYLENE INSENSTIVE 2 (EIN2), the essential positive regulator of ethylene signaling, at its C-terminal end (EIN2-C), leading to the repression of the EIN2 activity^17,18^. In the absence of ethylene, EIN2 is localized to the ER membrane, where it interacts with two F-box proteins, EIN2 TARGETING PROTEIN 1/2 (ETP1/2), mediating its protein degradation via the ubiquitin-proteasome pathway^19^. Upon the perception of ethylene, both ETR1 and CTR1 are inactivated, and the EIN2-C is dephosphorylated through an unknown mechanism^18^. The dephosphorylated EIN2-C is cleaved and translocated into both the nucleus and the P-body^18,20,21^. In the P-body, EIN2-C mediates the translational repression of two F-box proteins EIN3-BINDING F BOX PROTEIN 1/2 (EBF1/2) to promote the protein accumulation of ETHYLENE INSENSITIVE 3 (EIN3), the key transcription activator that is sufficient and necessary for activation of all ethylene-response genes^6,21,22^. In the nucleus, EIN2-C mediates the direct regulation of histone acetylation of H3K14 and H3K23 via histone binding protein EIN2 NUCLEAR ASSOCIATED PROTEIN 1/2 (ENAP1/2), leading to an EIN3 dependent transcriptional regulation for ethylene response^23–25^.

Epigenetic regulation in gene expression plays critical roles in various plant developmental processes and stress response^26–30^. Histone acetylation promotes the relaxation of nucleosomes between wrapped DNA and histone octamer by neutralizing the positive charges of lysine residues in histones, which is crucial for all DNA-based processes, including DNA replication and gene transcription^31,32^. Histone acetylation has been implicated in plant response to different hormones including ABA, Auxin, BR, JA and ethylene^33–35^. Several histone acetyltransferases (HATs) and histone deacetylases (HDACs) are implicated in the ethylene response. Plants with mutations in HAC1 and HAC5 (*hac1hac5*) displayed ethylene hypersensitive ethylene phenotype, and ethylene biosynthesis genes are upregulated in the double mutant, suggesting that HAC1 and HAC5 repress downstream ethylene responsive genes, lead to hyper ethylene responsive genes^36^. HDA6 is a member of histone deacetylase that interacts with both ethylene-stabilized master transcription factors, EIN3 and EIN3-LIKE 1 (EIL1), and JA-degraded JAZ to regulate JA-induced derepression of ethylene-responsive genes^36^, which mediates the crosstalk between ethylene and JA. HDA19, another member of HDAC, elevates the expression of target genes to trigger ethylene response^37^. Research from our group has found that NAD-dependent histone deacetylase SRT1 and SRT2 interact with ENAP1 to regulate subset of ethylene repressed genes in the ethylene response^38^.

A number of studies have focused on the role of histone acetylation in ethylene signaling, and our previous studies have shown that EIN2 directly regulates H3K14ac and H3K23ac in response to ethylene, which requires EIN3-mediated positive feedback regulation^39^. However, histone acetyltransferase involved in EIN2-directed histone acetylation remains unknown. In this study, we identified the histone acetyltransferase HAF2, which regulates H3K14ac and H3K23ac in response to ethylene. Our genetic and molecular findings demonstrated that mutations in HAF2 confer plants with ethylene insensitivity. We then discovered that HAF2 interacts with EIN2-C in response to ethylene. Biochemical assays revealed that the bromodomain of HAF2 binds to H3K14ac and H3K23ac peptides with a distinct preference for H3K14ac; the HAT domain possesses acetyltransferase catalytic activity for H3K14 and H3K23 acetylation, with a preference for H3K14. Our ChIP-seq data provide further evidence that HAF2 regulates H3K14ac and H3K23ac levels, and their ethylene-induced elevation is impaired in the *haf2* mutant. Moreover, our findings reveal that HAF2 co-functions with pyruvate dehydrogenase complex (PDC) to regulate H3K14ac and H3K23ac in response to ethylene in an EIN2 dependent manner. In summary, our research unveils the role of HAF2 as a histone acetyltransferase, forming a complex with PDC and EIN2-C to regulate histone acetylation of H3K14 and H3K23, preferentially for H3K14 in response to ethylene.

## Result

### HAF2 is involved in ethylene response

Our previous studies have shown that EIN2 directly regulates H3K14ac and H3K23ac in response to ethylene, a process that requires EIN3-mediated positive feedback regulation^39^. However, histone acetyltransferase involved in EIN2 mediated histone acetylation remains unknown. Here we found that mutations in *HAF2* conferred plants (*haf2-ko* and *haf2-del*) a partial ethylene-insensitive phenotype (Fig. 1A and 1B, Fig. S1A-1C). To further validate the function of HAF2 in ethylene response, we attempted to generate a gain-of function of HAF2 (*HAF2-Fox*), however, we did not obtain any plants that overexpressed *HAF2*, and the protein levels in *HAF2-Fox* were similar to those in native promoter driven *HAF2 (pHAF2-HAF2-GFP)* complementation lines (Fig. S1D). Subsequently we generated a truncated form of *HAF2-MC* and generated its overexpression plants (*HAF2-MCox*) (Fig. S1E). The resulting plants displayed a constitutive ethylene responsive phenotype even in the absence of ethylene (Fig. 1A and 1B), and the phenotype is more pronounced in the presence of ethylene, providing a genetic evidence supporting the involvement of HAF2 in the ethylene response.

**Figure 1.**
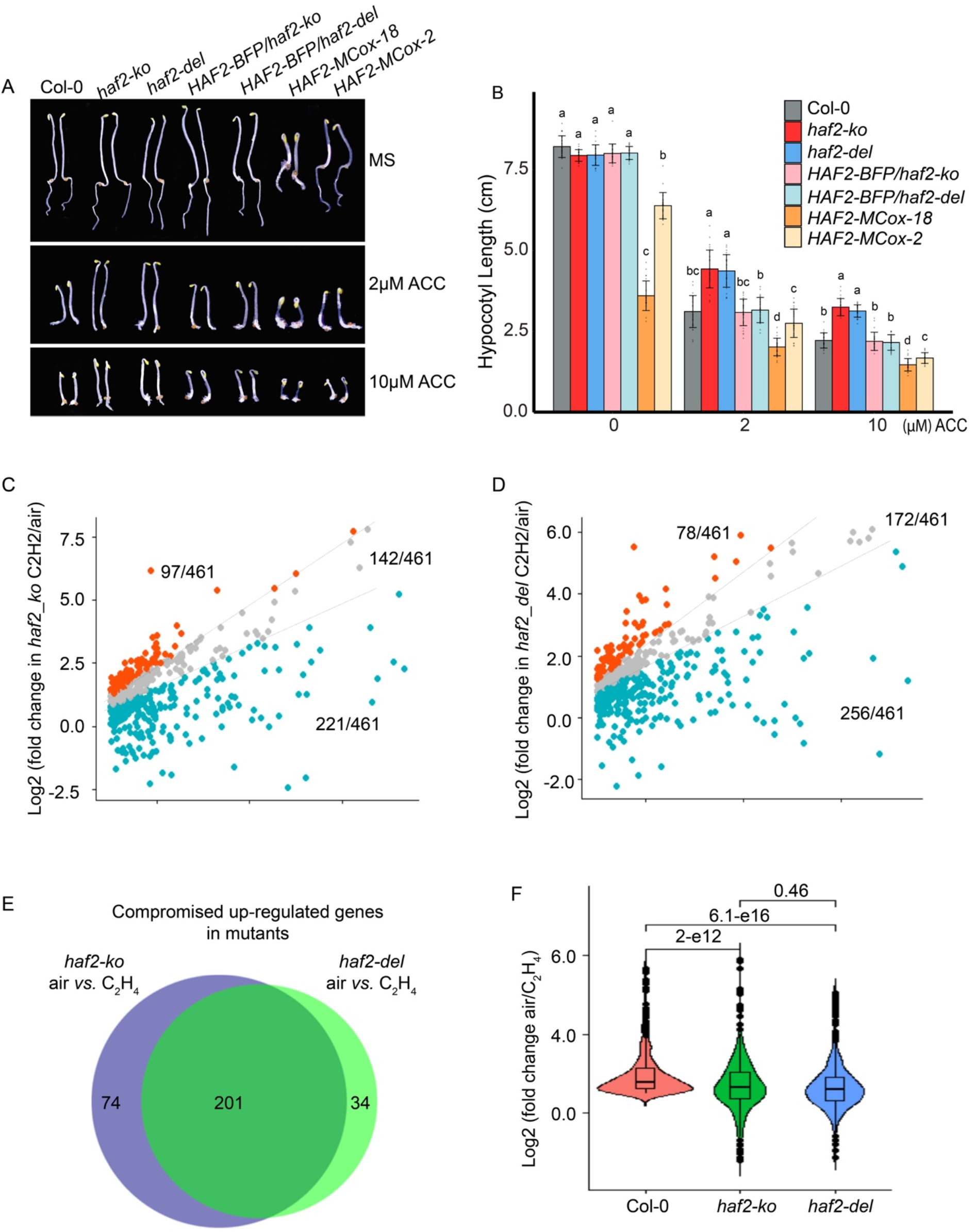
HAF2 is involved in ethylene response. (**A**) Photographs of seedlings with mutations in *haf2* mutants complementation, *HAF2MCox,* and Col-0 grown on MS medium containing 2 μM and 5 μM ACC with or without ACC. (**B**) Measurements of hypocotyl lengths of indicated mutants. Values are means ± SD of at least 20 seedlings. (**C** and **D**) Scatter plots to compare the log_2_(Fold Change) of ethylene up-regulated genes in Col-0 with that in *haf2-ko* (**C**) or in *haf2-del* (**D**) in mRNA-seq. Each dot represents a gene whose transcription level is significantly elevated by ethylene gas treatment in Col-0 (log_2_(Fold Change) > 1, *p*-adjust < 0.05). (**E**) Venn diagram to show the ethylene up-regulated genes that were compromised in *haf2-ko* and in *haf2-del*. (**F**) Violin plot of the distributions of log_2_(Fold Change) of ethylene up-regulated genes in Col-0 and in the indicated plants. *P* values were determined by a two-tailed *t* test.

To further evaluate the function of PDC in ethylene response at a molecular level, we performed mRNA sequencing (mRNA-seq) using *haf2-ko and haf2-del* seedlings treated with or without 4 hours of ethylene gas (Fig. S2A). We found that about 50% genes up-regulated by ethylene in Col-0 were not differentially expressed in either of the two mutants in response to ethylene (Fig. S2B). For further statistical quantification, we plotted the log_2_ fold change (log_2_FC) of each gene up-regulated by ethylene in Col-0 against that in each mutant. These genes were then divided into three groups: elevated group (log_2_FC in mutants is 30% greater than in Col-0), unchanged group (log_2_FC in mutants is within 30% of that in Col-0), and compromised group (log_2_FC in mutants is 30% less than that in Col-0). We found that the elevation of more than half of ethylene-up regulated genes in Col-0 was significantly compromised in both double mutants (Fig. 1C and 1D and Table S1-2). Importantly, most of the compromised genes identified from *haf2-ko* mutant seedlings were also compromised in the *haf2-del* seedlings (Fig. 1E), and the log_2_FC profile showed high similarity (Fig.1F). Further gene ontology analysis using those upregulation compromised genes showed that the ethylene signaling genes were overrepresented (Fig. S2C). Together, these data provide genetic and molecular evidence that HAF2 is involved in the ethylene response.

### HAF2 interacts with EIN2 via its C-terminal end

HAF2 contains an HAT domain, a ubiquitin binding domain in the middle region, a ZnF domain and a bromodomain at the C-terminal end (Fig. 2A). To further explore how HAF2 functions in the ethylene response, we truncated HAF2 as HAF2-N, HAF2-M and HAF2-C based on different domains (Fig. 2A), and examined their interactions with EIN2-C by yeast two-hybrid. We found that EIN2-C strongly interacted with HAF2-C, but not with HAF2-N end or HAF2-M. Notably, no interactions between EIN3 with any truncated HAF2s were detected (Fig. 2B). To further confirm the interaction, we conducted reciprocal in vitro pull down assay using in vitro purified proteins. Consistently, a strong physical interaction between EIN2-C and HAF2-C or HAF2-MC was detected, but no interaction was detected between EIN2-C and either HAF2-N or HAF2-M. This finding indicates that the C-terminal end of HAF2 is crucial in mediating its interaction with EIN2-C (Fig. 2C and 2D).

**Figure 2.**
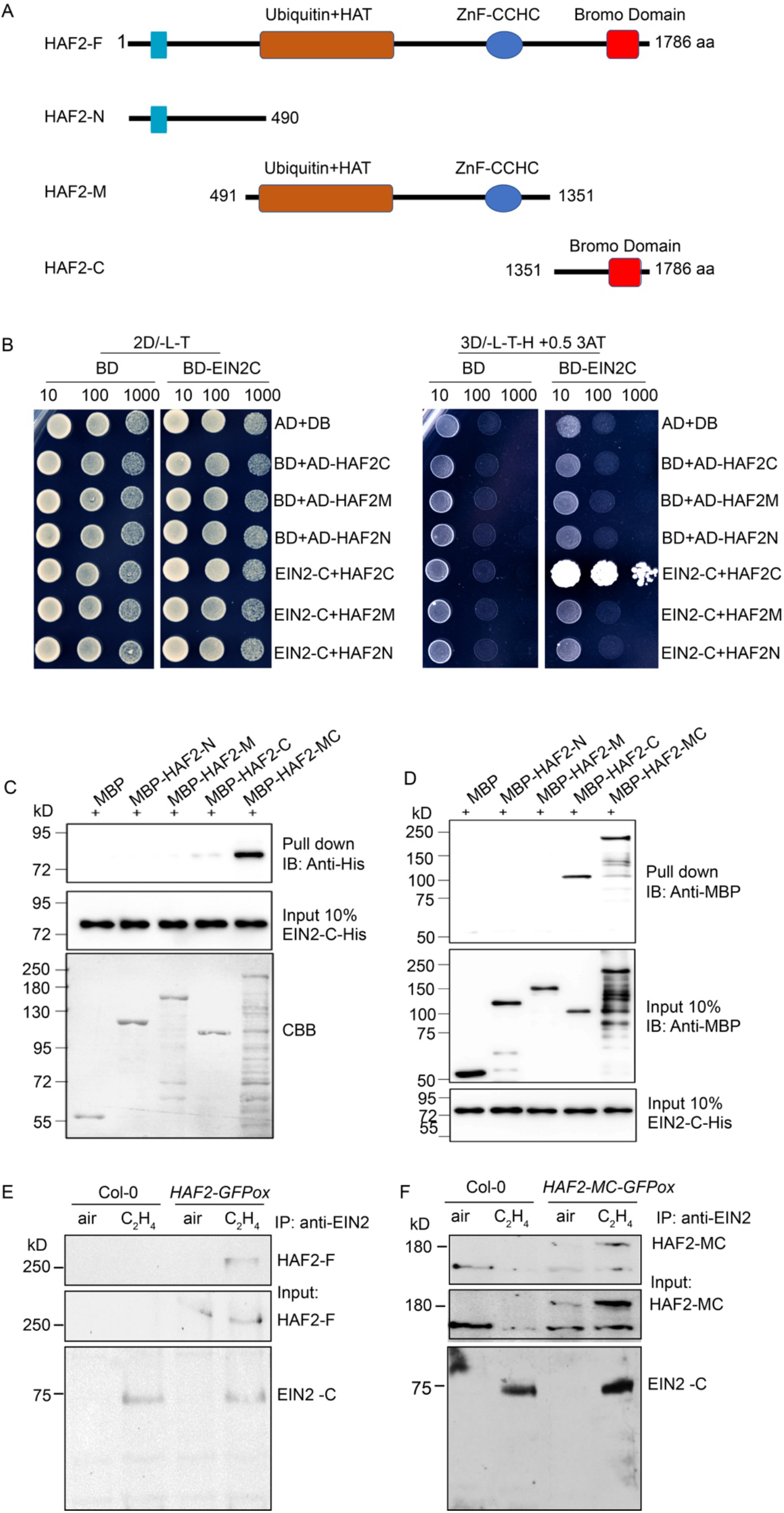
HAF2 interacts with EIN2-C via its C-terminal end. (**A**) Diagram to show the different truncated HAF2s that applied for the yeast two hybrid assay. (**B**) Yeast two-hybrid assay to detect the interaction between EIN2-C and HAF2-N, HAF2-M or HAF2-C. The indicated constructs were co-transformed into yeast. Yeasts were grown on the Leu and Trp drop-out medium(SD/-L-W) serving as the loading control (left panel). Yeasts were grown on Leu(L),Trp (W) and His(H) drop-out medium(SD/-L-W-H) with 3’AT to evaluate protein-protein interaction (right panel). **(C-D)** Pull down assay to show the interaction between EIN2-C and HAF2. MBP was used as negative control, MBP tagged HAF2-N, -M, -C and -MC were used as bait proteins (**C**), or EIN2-C-His was used as bait protein (**D**). (**E-F**) *In vivo* co-immunoprecipitation assay to examine the interaction between EIN2-C and HAF2 (**E**) or HAF2-MC (**F**) in the indicated transgenic plants. Total protein extractions from 3-day-old etiolated seedlings of indicated plants treated with or without 4 hours ethylene gas were applied for the assay.

To further confirm the interaction between EIN2-C and HAF2 in vivo, we performed in vivo co-immunoprecipitation (Co-IP) assays in *HAF2-GFP* and *HAF2-MC-GFP* etiolated seedlings treated with or without 4 hours of ethylene gas. As shown in Figures 2E and 2F, both HAF2 full length protein levels and HAF2-MC protein levels were upregulated by ethylene, and the interaction between EIN2 and HAF2 or HAF2-MC was detected in the presence of ethylene treatment. We further examined the cellular localization of HAF2-GFP and HAF2-MC-GFP in the presence of ethylene. The result showed that both HAF2-GFP and HAF2-MC-GFP were exclusively localized to the nucleus in the presence of ethylene (Fig. S3). All together, these results demonstrate that HAF2 and EIN2 interact in the presence of ethylene, and the C-terminus is important to mediate the interaction.

### The bromodomain of HAF2 binds to H3K14ac and H3K23ac

A bromodomain is a conserved structural module found in chromatin- and transcription-associated proteins that acts as the primary reader for acetylated lysine residues (Fig. S4A)^40,41^. AlphaFold prediction of HAF2 shows a four-helical bundle that appears to be arranged in a bromodomain-like motif^42^ (Fig. 3A). We designated the four helixes (from N to C) Z, A, B, and C, following the conventional labelling of other bromodomains (Fig. 3A). Additionally, like several other bromodomains, the C helix of HAF2 is predicted to be a long-bent helix^42,43^, segmented into a pair of helices that are approximately 34 Å and 24 Å long.

**Figure 3.**
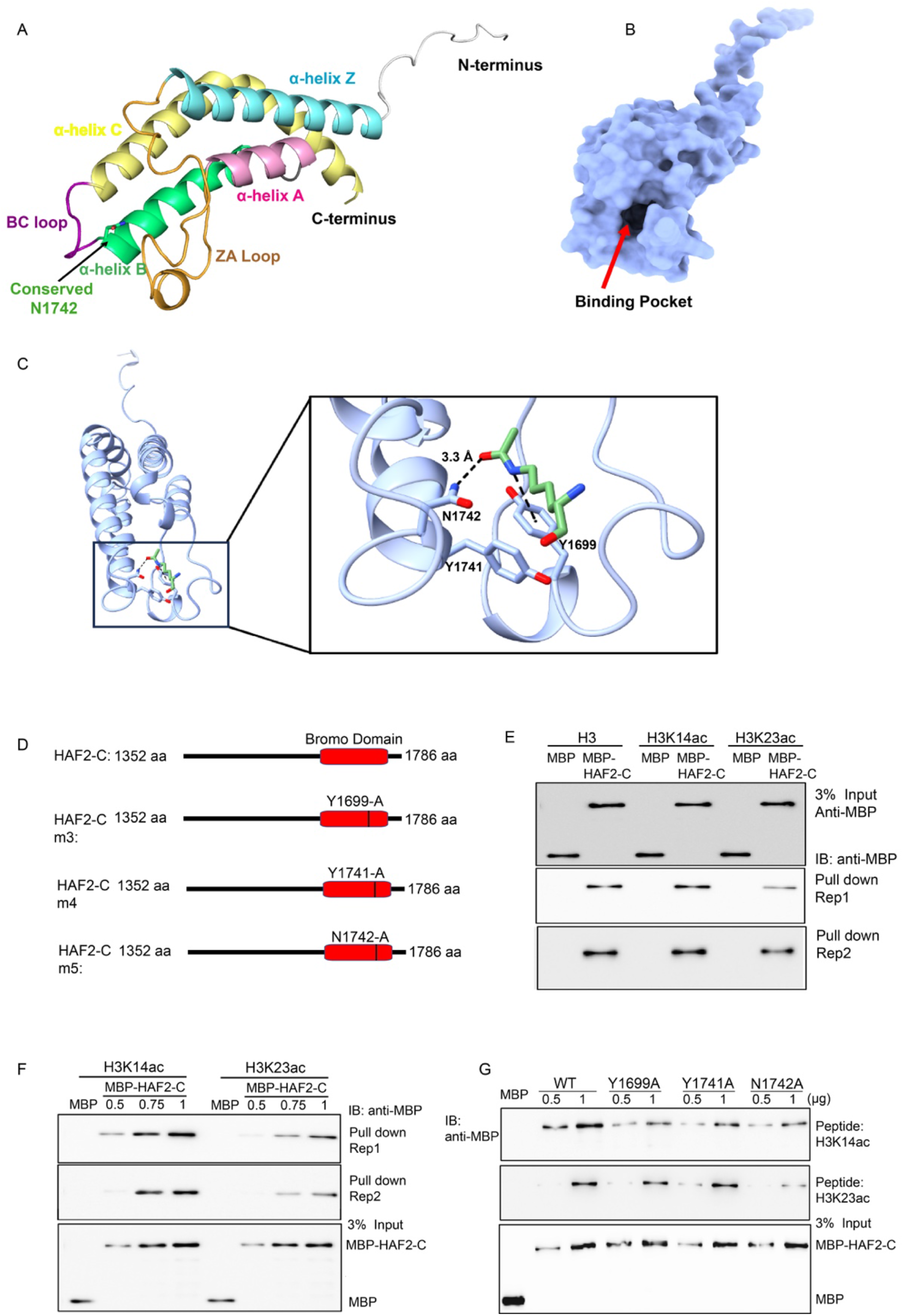
The bromodomain of HAF2 binds to H3K14ac and H3K23ac and the N1742 residue is important for the binding activity. (**A**) AlphaFold2 structure prediction of HAF2. The four helices Z, A, B, C are colored light blue, pink, green, and yellow, respectively. The ZA loop is colored orange, and the BC loop is colored purple. The conserved N1742 residue is shown as sticks. (**B**) Surface representation of the HAF2 AlphaFold2 model, where the deep acetyl-lysine binding pocket is visible at the center of the structure. (**C**) Model of an acetyl-lysine residue bound to HAF2. The acetyl-lysine residue is colored green. HAF2 is colored light blue, with the key residues Y1699, Y1741, and N1742 shown as sticks. Distance measurements representing HAF2-ligand interactions are shown as black dashed lines. (**D**) Diagrams to show the C-terminus of HAF2 that contains wild type bromodomain and mutated bromodomains. (**E**) In vitro histone peptide binding assay using in vitro purified recombinant wild type HAF2-C. MBP protein served as a negative control. (**F**) In vitro histone peptide binding assay to detect the binding activity with different concentrations of HAF2-C. (**G**) In vitro binding assay to show the impact of indicate amino acids on the histone peptide binding activity of HAF2 bromodomain. The proteins applied for the binding assays were purified from the in vitro expression system.

To understand how the HAF2 bromodomain recognizes histone substrates, we compared the predicted HAF2 structure to other published bromodomain structures in complex with histone peptides. Examination of these structures (PDB code: 2RNW^44^, 6KMJ^45^, 3O34^46^) revealed that the acetylated lysine binding region is consistently located at the end of the helical bundle, close to the ZA and BC loops (Fig. 3B). Despite the similar placement of the acetyl-lysine residue within the binding pocket of published bromodomain complex structures, the bound histone peptides adopt different configurations in their polypeptide backbone upon peripherally associating with a given bromodomain. Furthermore, the peripherally associated histone peptides exhibit relatively limited interactions with the bromodomain. An exception to this trend is the notable interactions observed between the bromodomain and the acetyl-lysine. The binding site of the acetyl-lysine residue is highly conserved in all bromodomain structures with a deep pocket (of approximately 11.6 Å) formed between the B helix and the ZA loop (Fig. 3C)^43^. This pocket is also where the specific inhibitor JQ1 binds to abolish the interaction between acetyl-lysine and the bromodomain^47^ (Fig. S4B and S4C). Thus, this deep pocket is also likely the site for acetyl-lysine recognition by HAF2. When we conducted modeling using MASTREO, we noticed that the highly conserved Asn1742 is in position to form a hydrogen bond with the acetyl group carbonyl (Fig. 3C). The formation of an amide bond between lysine and the acetyl group adapts a sp^2^ hybridization, stacking in a parallel manner to the aromatic ring of Tyr1699. Furthermore, Tyr1741 forms a perpendicular π-π stacking interaction, greatly stabilizing the interaction of the substrate acetyl-lysine with Tyr1699.

To further examine whether the bromodomain in HAF2-C binds to H3K14ac or H3H23ac, we performed a histone peptide binding assay using in vitro purified HAF2-C that includes bromodomain. The result showed that HAF2 bound to both H3K14ac and H3K23ac peptides; however, the HAF2-C exhibited a stronger preference for binding to the H3K14ac peptide compared to H3K23ac (Fig. 3D-3F). To further assess whether the binding activity of HAF2-C is conferred by its bromodomain, we introduced mutations to the residues (Tyr1699, Tyr1741, Asn1742) predicated as crucial for bromodomain binding activity, as depicted in Figure 3A-C. We created mutated variations of HAF2-C that carrying on single point mutations in residues of Y1699, (1699 Y-A, HAF2-C3) ,Y1741 (1714 Y-A, HAF2-C4) and Y1742 (1742 N-A, HAF2-C5), respectively (Fig. 3D). Subsequent binding assays using those mutated proteins revealed that the mutation of Tyr1699 reduced the association of HAF2-C to H3K14ac peptide but not to H3K23ac peptide (Fig. 3G). Similarly, a mutation of Tyr1741 only reduced association with the H3K14ac histone peptide. However, a mutation of Asn1742 substantially reduced association with both histone peptides (Fig. 3G). These results reveal that the bromodomain in HAF2 exhibits binding activity to H3K14ac and H3K23ac; the residues Tyr1699, Tyr1741 regulate the binding of the HAF2 bromodomain to H3K14ac, whereas the residues N1742 is identified as a key residue for the binding activity to both H3K14ac and H3K23ac. This is explained by the structural model, because a hydrogen bond between Asn1742 and the acetyl group of the substrate is a key interaction for acetyl-lysine recognition (Fig. 3C).

As we showed that overexpression of HAF2-MC leads to a hypersensitive ethylene response. To further evaluate whether the mutations in bromodomain will impact the function of HAF2 in ethylene response, we generated plants overexpressing *HAF2-MC3 (HAF2-MC3ox),* and *HAF2-MC5* (*HAF2-MC5ox).* We have screened over 50 individual transgenic with either *HAF2-MC3ox* or *HAF2-MC5ox.* Among them, three individual *HAF2-MC3ox* plants displayed a mild ethylene hypersensitivity, whereas, two of *HAF2-MC5ox* plants displayed a very minor ethylene hypersensitive phenotype (Fig. S4D and 4E). This provides genetic evidence that the bromodomain is important for the functions of HAF2 in ethylene response, and N1742 is a crucial residue for this function.

### HAF2 is a histone acetyltransferase that regulates the acetylation of H3K14ac and H3K23ac both in vitro and in vivo

HAF2 is annotated as histone acetyltransferase. A previous study showed that mutated *HAF2* resulted in a decrease of H3ac in target genes by ChIP-PCR^48^. However, there is currently no available biochemical data confirming its enzymatic activity. Given that the acetylation of histone at H3K14 and H3K23 is induced by ethylene, we decided to examine whether the HAT domain in HAF2 has histone acetyltransferase activity for H3K14 and H3K23 acetylation. To achieve this goal, we constructed proteins of HAF2-M that contains the HAT domain, HAF2-MC that contains both the HAT domain and the bromodomain, and HAF2-C that contains only the bromodomain. We then examined their histone acetyltransferase (HAT) activity by in vitro enzyme activity assay using in vitro purified proteins with mono nucleosomes or in vitro purified histone H3 as substrates. Weak enzyme activity was detected for both H3K14 and H3K23 acetylation from HAF2-M, but not from HAF2-C (Fig. 4A and 4B). But, the enzyme activity was significantly enhanced when the C-terminal domain was present. To further evaluate the functions of the bromodomain in the enzyme activity, we constructed the HAF2-MC-KO, carrying on the same mutations as that in the *haf2-ko* mutant, in which the bromodomain was abolished and the plant showed partial ethylene insensitive phenotype (Fig. 1A and Fig. S1A, S1B). We then performed an enzyme activity assay. The result showed that the HAT activity for H3K14 was still detectable from HAF2-MC-KO using histone H3 as a substrate, but was substantially reduced compared to the wild type HAF2-MC (Fig. 4A). While no HAT enzyme activity for H3K23ac was detected from HAF2-MC-KO either with mono nucleosomes or histone H3 as substrates (Fig. 4B), providing further evidence showing that HAF2 bromodomain plays a pivotal role in the regulation of HAF2 HAT activity. Altogether, these results demonstrate that HAF2 functions as a histone acetyltransferase that is capable of acetylating H3K14 and H3K23, the bromodomain plays a critical role in regulating the enzyme activity.

**Figure 4.**
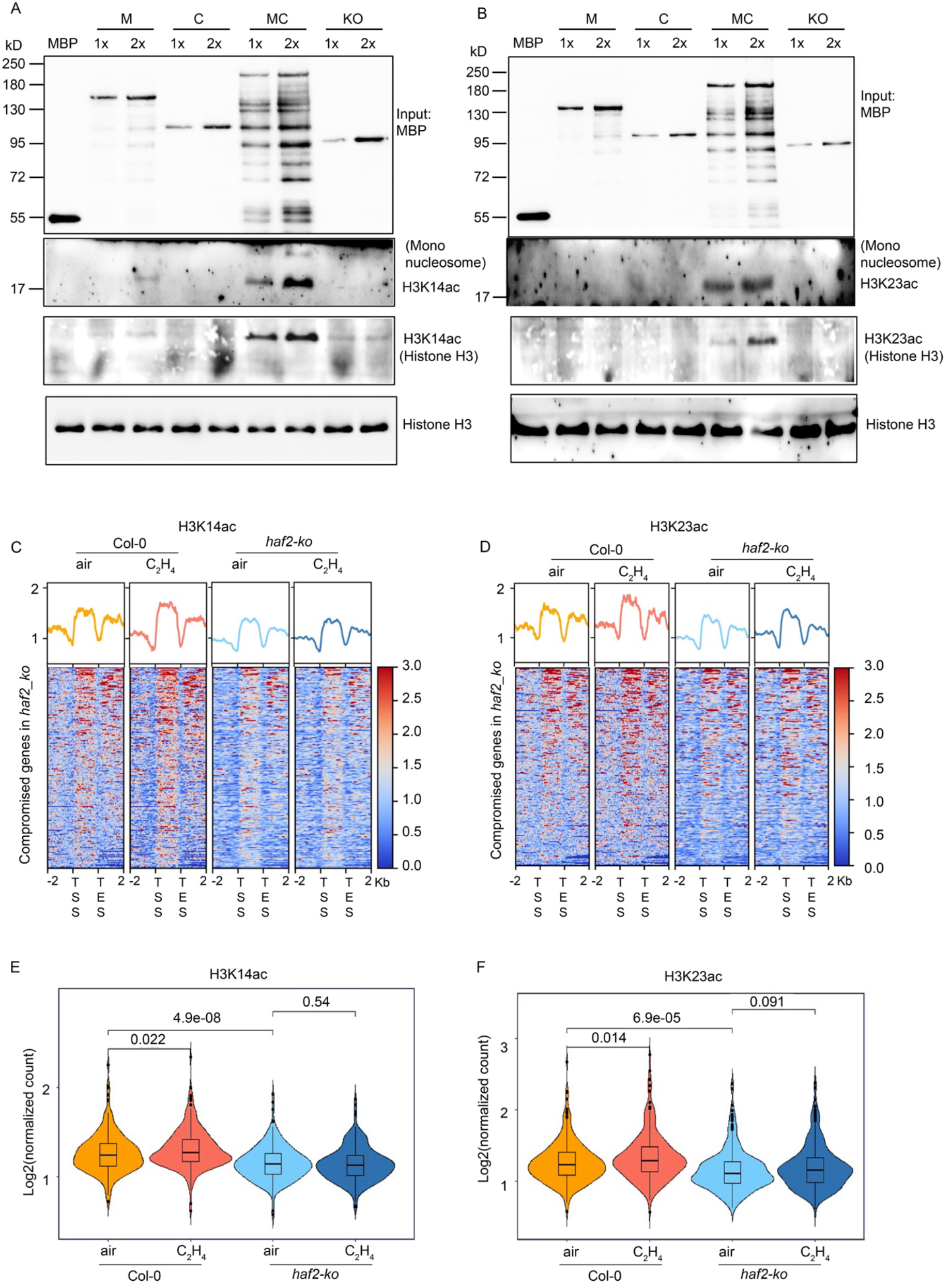
The HAT domain of HAF2 has histone acetyltransferase activity for H3K14 and H3K23. (**A-B**) In vitro histone acetyltransferase activity assay of the recombinant proteins indicated in the figure. Human mononucleosomes or in vitro purified histone H3 served as substrates, the acetylated histone H3 were detected by anti-H3K14ac (**A**) and anti-H3K23ac (**B**) antibodies. **(C** and **D**) Heatmaps of H3K14ac (**C**) and H3K23ac (**D**) ChIP-seq signals (log_2_ ChIP signal) from the genes whose ethylene-induced expressions are co-compromised in the *haf2-ko* and the *haf2-del* mutants. TSS stands for transcription start site; TTS stands for transcription termination site. (**E and F**) Violin plots illustrating log_2_ normalized H3K14ac ChIP signals (**E**) and H3K23ac ChIP signals (**F**) from TSS to TES that in the genes that were co-compromised in the *haf2-ko* and the *haf2-del* mutants. *P* values were calculated by a two-tailed *t* test.

To investigate the regulations of HAF2 on H3K14ac and H3K23ac in response to ethylene in vivo, we conducted chromatin immunoprecipitation sequencing (ChIP-seq) for H3K14ac and H3K23ac in the *haf2-del* mutant treated with or without 4 hours of ethylene. We observed a reduction in H3K14ac and H3K23ac levels in the *haf2-ko* mutant compared to Col-0 over genes with a compromised transcriptional activity in the *haf2-ko* mutant. Furthermore, the ethylene-induced elevation of H3K14ac and H3K23ac was impaired in the *haf2-ko* mutant (Fig. 4C, 4D, and Fig. S5A-C). We then evaluated the ChIP signals from TSS to TTSs. Similarly, a significant reduction was detected in the *haf2-del* mutant both with and without ethylene treatments, and the ethylene induced elevation was impaired in the *haf2-del* mutant (Fig. 4E and 4F). We subsequently conducted H3K14ac and H3K23ac ChIP-qPCR assays on selected target genes in Col-0, the *haf2-ko* mutant and *HAF2-MCox,* treated with and without 4 hours of ethylene. The results from Col-0, the *haf2-del* mutant further validated the ChIP-seq data. Furthermore, the H3K14ac levels were significantly elevated in *HAF2-MCox,* even in the absence of ethylene, whereas, only a slight elevation of H3K23ac was detected in *HAF2-MCox* in the absence of ethylene (Fig. S5B and S5C), indicating the preference of HAF2 for H3K14ac.

To further evaluate the impact of histone acetylation regulation on gene expression, we conducted qRT-PCR assays. The results showed that the expression of target genes was positively correlated with the levels of H3K14ac and H3K23ac. The ethylene-induced activation of gene expression was impaired in the *haf2* mutants. Whereas, in *HAF2-MCox*, gene expression was elevated even in the absence of ethylene, and ethylene induced a more significant elevation (Fig. S5D). Collectively, these results demonstrate that HAF2 functions a histone acetyltransferase, acetylating H3K14 and H3K23 to facilitate active gene expression in response to ethylene, with a preference for H3K14ac.

### HAF2 and pyruvate dehydrogenase complex (PDC) co-function to regulate histone acetylation in response to ethylene

We have found the PDC complex can translocate to the nucleus to interact with EIN2-C to provide acetyl CoA for the regulation of H3K14ac and H3K23ac in response to ethylene^49^. Given HAF2 is shown to regulate H3K14ac and H3K23ac in the ethylene response. we decided to examine whether HAF2 and PDC function in the same pathway. We first examined the interaction between PDC and the HAF2-N, HAF2-M or HAF2-C by a semi in vitro pull down assay. However, no interaction was detected. We further compared the *haf2-del* transcriptome with that of *e1-2-2e2-2* in response to ethylene. The transcriptomes profile of these two mutants in response to ethylene were very similar, and the majority of the compromised ethylene-activated genes in these two mutants were overlapped (Fig. 5A and 5B). Gene ontology analysis using those upregulation compromised genes in both the *haf2* mutants and the *e1-2-2e2-17* mutants showed that the ethylene signaling genes were overrepresented (Fig. 5C), suggesting that HAF2 and PDC share a similar regulation in the ethylene response at a molecular level. Hence, we decided to explore their genetic connections by generating *haf2e1e2* triple mutants. Due to the linkage of *HAF2* and *PDC-E2,* we created *HAF2* mutations in the *e1-2-2e2-17* plants (*haf2-3 e1-2-2e2-17* and *haf2-49 e1-2-2e2-17*) by CRISPR-Cas9 (Fig. S6A). Notably, the mutations in *haf2-3 e1-2-2e2-17* and *haf2-49 e1-2-2e2-2* closely resembled those in the *haf2-ko* mutant (Fig. S1 and Fig. S6). We then compared the ethylene responsive phenotype of the *haf2-del* single mutant with that of *haf2-3e1-2-2e2-17* and *haf2-49e1-2-2e2-17* mutants. The result showed that the ethylene insensitivity of *haf2-del* or *e1-2-2e2-2* was enhanced in both *haf2-3e1-2-2e2-17* and *haf2-49e1-2-2e2-17* (Fig. 5D and 5E).

**Figure 5.**
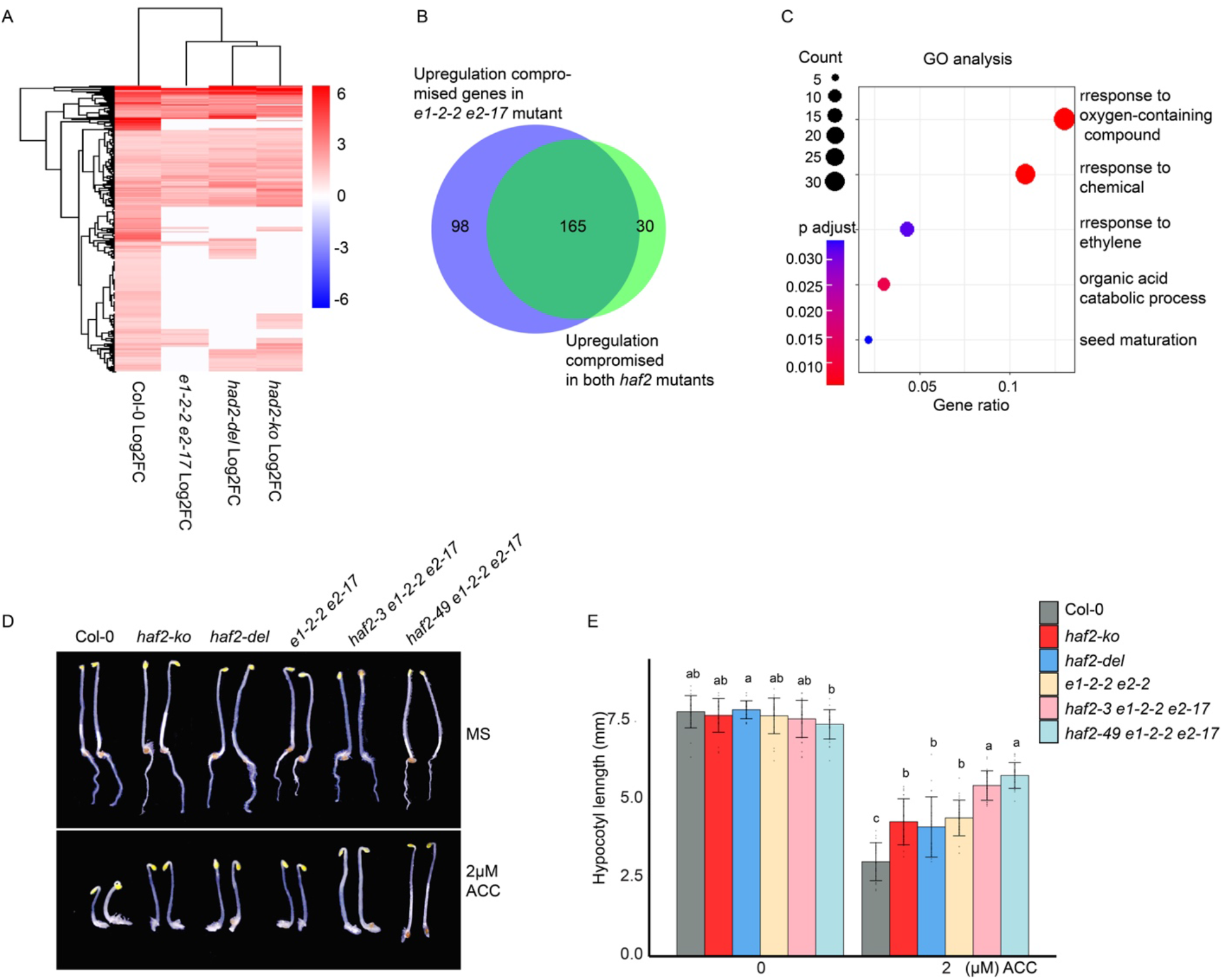
HAF2 co-functions with PDC to regulate ethylene response. (**A**) Heatmap of log_2_(Fold Change C_2_H_4_/Air) of all ethylene up-regulated differential expressed genes that were identified in Col-0 in *e1-2-2 e2-17* and in the *haf2* mutants. Raw log_2_FC value of each gene is annotated in the heatmap. (**B**) Venn diagram to show the overlap of compromised ethylene up-regulated genes in *e1-2-2 e2-17* and in the *haf2* mutants. (**C**) GO analysis of the compromised ethylene up-regulated in both *e1-2-2 e2-17* and the *haf2* mutants. (**D**) Photographs of seedlings of indicated plants on MS medium containing 2 μM ACC or without ACC. Plants were grown in the dark for three days before beings photographed. (**E**) Measurements of hypocotyl lengths of indicated mutants. Values are means ± SD of at least 20 seedlings. different letters indicate statistically significant differences (One-way ANOVA test followed by Tukey’s HSD test, *P* ≤ 0.05).

As both PDC and HAF2 are involved in the regulation of histone acetylation in response to ethylene, we conducted a comparative analysis of ChIP-seq profiles for H3K14ac and H3K23ac between the *haf2-ko* and the *e1-2-2e2-2* mutants in response to ethylene. The result showed that the regulations of H3K14ac and H3K23ac by HAF2 and PDC were highly similar (Fig. 6A-6D). To further assess the impact of HAF2 and PDC on H3K14ac and H3K23ac, we performed H3K14ac and H3K23ac ChIP-qPCR assays in the selected targets. Our result showed that the ethylene-induced elevation of H3K14ac and H3K23ac was reduced both in the *haf2-3* mutant and in the *e1-2-2e2-17* mutant. Furthermore, this reduction was enhanced in both the *haf2-3e1-2-2e2-17* and the *haf2-49e1-2-2e2-17* triple mutants (Fig. 6E and 6F). We then examined the target gene expression by qRT-PCR. The ethylene-induced upregulation of the target gene expression was reduced, with a more significant reduction in both the triple mutants than in either the *e1-2-2e2-2* mutant or in *haf2* single mutants (Fig. 6G). To assess the potential collaborative function of HAF2 and PDC at the chromatin level, we conducted ChIP-qPCR of PDC-E2 and HAF2 in the *E2-GFP* and *HAF2-BFP* plants treated with air or 4 hours ethylene in selected target loci. The signal from both PDC-E2 and HAF2 binding was enriched in the promoters of the selected target genes (Fig. 7A-7C). However, in the absence of EIN2, the ethylene-induced elevated binding of PDC-E2 or HAF2 was eliminated, indicating that EIN2 is required for the ethylene-induced binding of PDC-E2 and HAF2 to the targets. Subsequently, we conducted a ChIP-qPCR assay, the results showed that HAF2 binding to the selected targets was EIN2 dependent (Fig. 7B). Taken all together, these results demonstrated that HAF2 and PDC function in the same complex to regulate H3K14ac and H3K23ac in response to ethylene, and HAF2 plays its roles in the ethylene-regulated genes depends on EIN2 (Fig. 7D).

**Figure 6.**
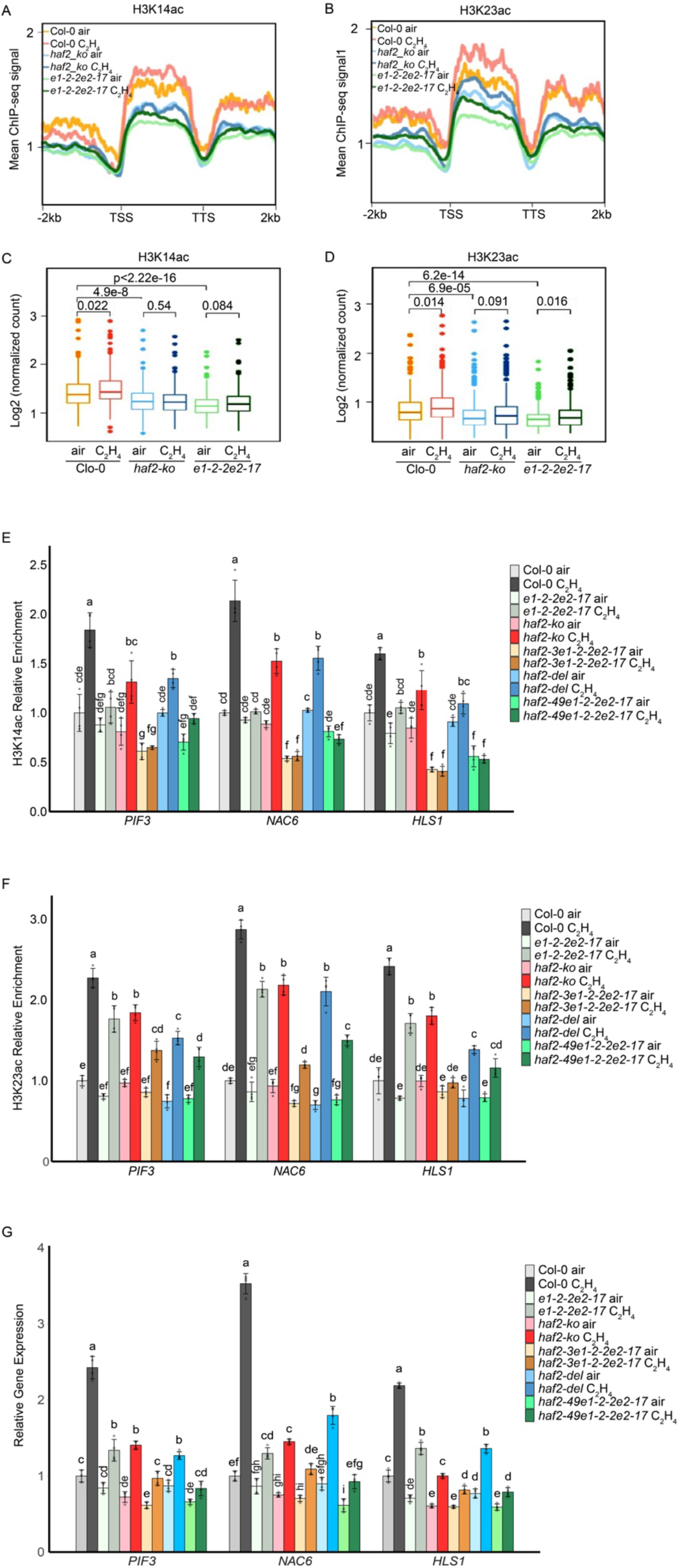
HAF2 cofunctions with PDC to regulate histone acetylation in response to ethylene. (**A-B**) Comparison of histone acetylation of H3K14ac (A) and H3K23ac (**B**) over ethylene up regulated genes in *haf2-ko* and *e1-2-2 e2-17* mutant. TSS stands for transcription start site; TTS stands for transcription termination site. (C-D) Box plots illustrating log_2_ normalized H3K14ac ChIP signal (**C**) and H3K23ac ChIP signal (**D**) from TSS to TTS that in the genes that were co-compromised in the *pdc e1-2-2 e2-17* double mutant in the indicated genotypes and conditions. *P* values were calculated by a two-tailed *t* test. (E-F) ChIP-qPCR to validate the enrichment of H3K14ac (**E**) and H3K23ac (**F**) in the indicated etiolated seedlings with or without 4 hours of ethylene treatment over representative genes. Individual data point of the relative fold change normalized to Col-0 air is plotted as a dot. Different letters indicate significant differences (*P* ≤0.05) between each genetic background and treatment condition calculated by a One-way ANOVA test followed by Tukey’s HSD test. (**G**) qRT-PCR analysis of target gene expressions in the indicated 3-day-old etiolated seedlings with or without 4 hours of ethylene gas treatment. Values are the average of relative expression from each gene normalized to that of *Actin2.* Error bars indicate the SD (n = 3) and different letters indicate statistically significant differences (One-way ANOVA test followed by Tukey’s HSD test, *P* ≤ 0.05).

**Figure 7.**
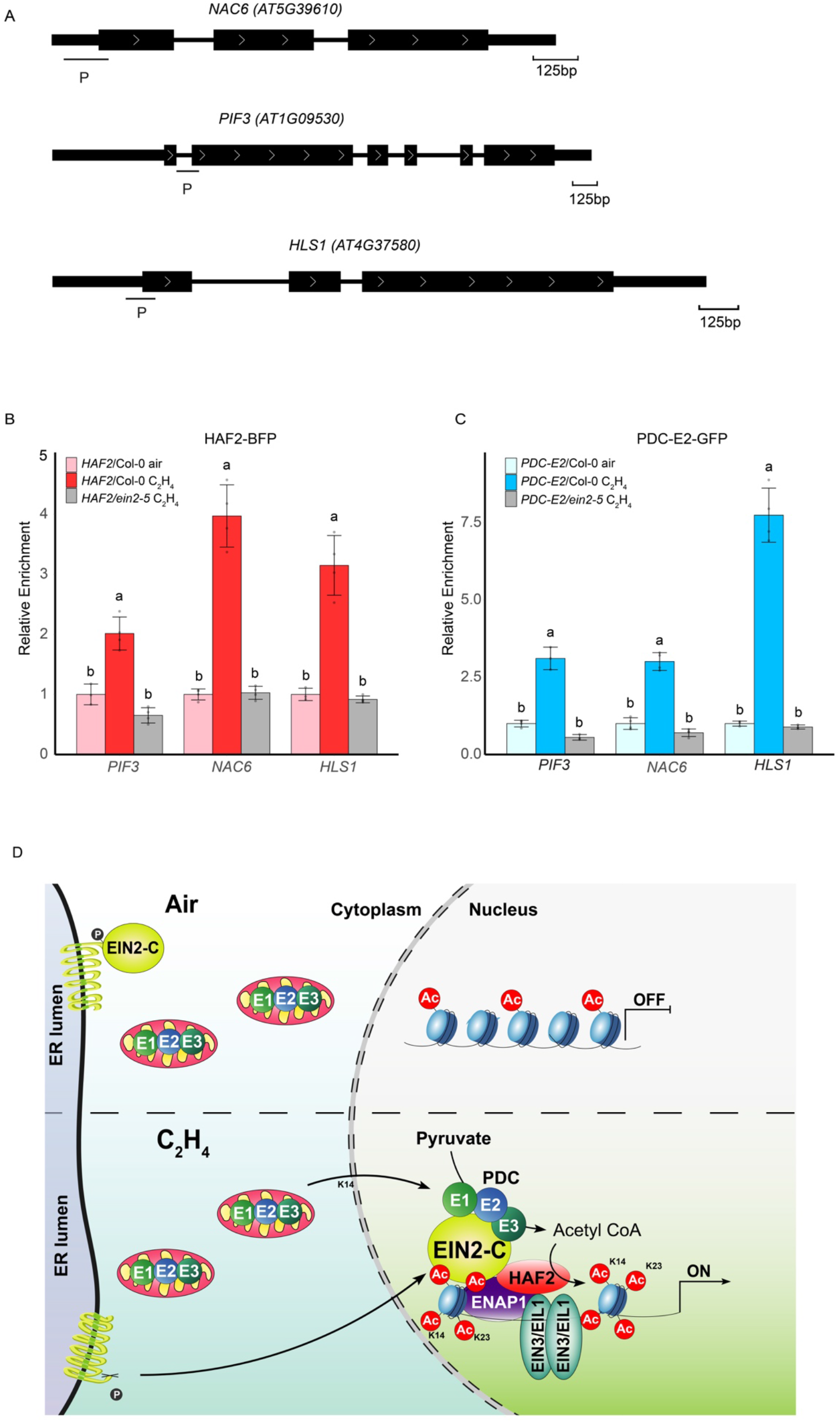
HAF2 and PDC binding to the same target in an EIN2 dependent manner. (**A**) Diagram to show the locations of primers for HAF2 binding and PDC-E2 binding assays. P indicates the binding target locus for ChIP-qPCR. (**B-C**) ChIP-qPCR to validate the enrichment of HAF2 (**B**) and PDC-E2 (**C**) in the indicated etiolated seedlings with or without 4 hours of ethylene treatment over representative genes. Individual data point of the relative fold change normalized to Col-0 air is plotted as a dot. Different letters indicate significant differences (*P* ≤0.05) between each genetic background and treatment condition calculated by a One-way ANOVA test followed by Tukey’s HSD test. (**D**) Model to illustrate HAF2’s function in ethylene response. In the absence of ethylene, there is no elevation of H3K14ac and H3K23ac (top). Upon the presence of ethylene, HAF2 interacts with EIN2-C and forms a complex with PDC to elevate histone acetylation at H3K14 and H3K23, thereby promoting EIN3-dependent transcriptional activation in response to ethylene (bottom).

## Discussion

Histone acetyltransferases (HATs) have been shown to play in various biological events, including the plant hormone responses^50–55^. Previously, we discovered that EIN2 mediates elevation of noncanonical histone acetylation at H3K14 and H3K23 elevation for transcriptional regulation in response to ethylene ^24^. Interestingly, we also observed that although H3K9ac levels are not regulated by ethylene, their levels in ethylene-upregulated genes are greater than those in ethylene-downregulated genes even in the absence of ethylene treatment ^35^. Further studies showed that SRT1 and SRT2 are two histone deacetylase that keep a low level of H3K9ac of a cluster of ethylene-repressed genes^38^. Despite the progress in understanding histone acetylation in the ethylene response, the histone acetyltransferase that directly functions with EIN2 for the regulation of H3K14ac and H3K23ac remains mysterious. In this study, we provide compelling evidence showing that HAF2 interacts with EIN2 and PDC to regulate histone acetylation H3K14ac and H3K23ac for transcriptional regulation in response to ethylene. Firstly, both our genetic and RAN-seq results clearly show that HAF2 regulates ethylene upregulated genes. Secondly, in vitro and in vivo assays support that HAF2 interacts with EIN2-C via C-terminal end. Third, we found that the bromodomain of HAF2 binds to H3K14ac and H3K23ac with a preference for H3K14ac. We further revealed that N1742 is one of the most important residues for the binding activity (Fig. 3). Fourth, our biochemical assay results demonstrate that the HAT domain has histone acetyltransferase activity, and the bromodomain can enhance the HAT activity (Fig. 4). Additionally, our ChIP-seq data provide a strong support for the biochemical assay results (Fig. 4). Finally, our genetic and molecular results show that HAF2 and PDC function together to regulate H3K14ac and H3K23ac in an EIN2 dependent manner in response to ethylene (Fig. 5 and 6).

In total, 12 HAT proteins were predicted in Arabidopsis, categorized into three distinct families including the GNAT-MYST family, P300/CREB-binding protein co-activator family and TAF_II_250 family. HAF2 belongs to TAF_II_250 family^56^. In addition to HAT domain, the HAF2 from Arabidopsis possess a single bromodomain, a zinc-finger-type C2HC domain is located at an approximately equal distance downstream of the HAT domain and a conserved ubiquitin signature at the N-terminal side of the HAT domain. A previous study indicated that HAF2 is required for light signaling, potentially through regulating histone H3 acetylation^48^. Later research showed that HAF2 targets to the circadian clock gene RRR5 and LUX to regulate H3ac for their gene expression^57^. However, no data has shown that HAF2 possesses histone acetyltransferase catalytic activity. In our study, we present biochemical evidence for the first time, demonstrating that the HAT domain within HAF2 processes histone acetyltransferase catalytic activity, targeting acetylation on H3K14 and H3K23 (Fig. 4). Significantly, the acetylation activity on H3K14 is greater than that on H3K23. Our binding assay results clearly demonstrate the HAF2 bromodomain’s preference for binding H3K14ac over H3K23ac (Fig. 3). Moreover, our data indicates that the bromodomain’s binding activity significantly enhances histone acetyltransferase (HAT) catalytic activity (Fig. 4A and 4B). Bromodomains are highly conserved 110–amino acid motifs that recognize acetyl-lysine residues^41^. Bromodomains have been identified in a variety of proteins, including transcriptional coactivators^58–60^, methyltransferases^61,62^, and HATs^63^. Zeng and Zhou defined that bromodomain can specifically recognize the acetylated lysine (KAc) mark on histones^64^. Human TFIID contains two tandem bromodomains, and the crystal structure of the double bromodomain reveals two side-by-side, four-helix bundles with a highly polarized surface charge distribution. Each bundle contains an Nɛ -acetyllysine binding pocket at its center, which results in a structure ideally suited for recognition of diacetylated histone H4 tails^59^.

Bromodomains share a conserved specialized structure that consists of four alpha-helices (αZ, αA, αB, and αC), linked by variable loop regions (ZA loop and BC loop), forming the largely hydrophobic KAc binding pocket. Alpha fold modeling result predict that the HAF2 bromodomain binding pocket can potentially bind H3K14ac and H3K23ac, and the N1742 is a vital residue for the binding activity to both H3K14ac and H3K23ac (Fig. 3). Our binding assays with wild type HAF2 bromodomain and mutated bromodomain further confirmed the modeling result. However, why the bromodomain has binding preference is unclear. Furthermore, in the enzyme activity assay, our findings indicate that the bromodomain has the ability to enhance the activity of the HAF2 HAT enzyme, displaying a preference for H3K14ac. This preference is likely linked to the binding specificity of the bromodomain. Therefore, acquiring the crystal structure of the HAF2 bromodomain and the combined structure of bromodomain and HAT domain is crucial to elucidate the underlying preference mechanisms. Many studies over the past decades have provided many insights into how the specificity of histone tail is determined for acetylation. However, the molecular mechanisms still remain unclear. Histone modification is orchestrated by a protein complex, and in addition to the histone modification reader, cofactors may also play a significant role in shaping specificity. Hence, identifying the composition of the protein complex and elucidating its structural details holds the promise of providing crucial and illuminating insights into this complex process.

Histone acetyltransferases catalyze acetylation of their cognate protein substrates using acetyl-CoA (Ac-CoA) as a cofactor. In mammal, three main cytosolic enzymes that synthesize acetyl CoA, including PDC, Acyl-CoA Synthetase Short Chain Family Member (ACSS2) and ATP Citrate Lyase (ACLY), have been shown to localize to the nucleus for various biological functions^65–71^. The PDC complex was reported to localize to the nucleus to interact with P300 in cell division to regulate cell proliferation^67^. ACSS2 catalyzes the conversion of acetyl CoA from acetate in the nucleus for the regulation of long-term spatial memory and glucose starvation response^68,69^. Nuclear ATP citrate lyase (ACLY) utilizes citrate to produce acetyl CoA to promote histone acetylation at double strand break sites for DNA repair^70^. More recently, a study showed that ACS2, the yeast ortholog of ACSS2, can be recruited to chromatin during quiescence exit and ACS2 is preferentially associated with the most up-regulated genes, suggesting that acetyl group transfer plays an important role in gene activation^72^. Studies in rice revealed that ACL translocates to the nucleus to interact with HAT1, and specifically acetylates H4K5 during cell division in developing endosperm^73^. Our research indicate that PDC can tanslocate to the nucleus to regulate H3K14ac and H3K23ac in response to ethylene^49^. Genetically and molecularly, HAF2 and PDC can enhance each other’s activity (Fig. 5 and 6). Our data also demonstrates that PDC and the HAF2 co-target the chromatin at selected loci (Fig. 7A and 7B), further supporting our previous study that PDC can tanslocates to the nucleus to regulate histone acetylation at H3K14 and H3K23 for transcription regulation in response to ethylene^49^. Further study on how PDC is translocated to the nucleus and how the PDC complex regulates HAF2 functions in response to ethylene will be a focus of future research.

## Materials and methods

### Plant materials, growth condition, and hormone treatments

*Arabidopsis* (*Arabidopsis thaliana*) accession Columbia (Col-0) was used as the wild type in this study and all the mutant genotypes are in this ecotype background. *Arabidopsis* seeds were first surface-sterilized with solution containing 50% bleach and 0.01% Triton X-100 for 10 minutes, washed with sterile distilled water for four times, sown on Murashige and Skoog (MS) medium plates containing 0.2% sucrose and 1% phytoblend agar and stratified for 3 days at 4 °C in the dark. For seed propagation, one- or two-week-old green seedlings were transferred into soil (Promix-HP) and grown in a growth chamber setting with the long-day photoperiod (16 h light/8 h dark) at 22 °C till maturity.

Ethylene triple response assay was performed with sterilized seeds on MS plates containing various concentrations of 1-aminocyclopropane-1-carboxylic acid (ACC, Sigma) (0, 1 μM, 2 μM, 5 μM, and 10 μM ACC). Approximately 60 seeds from each genotype collected in the similar time period were sown on the same MS medium plate per concentration. Seeds sown on ACC plates were kept at 4 °C in the dark for 3 days for stratification and then they were exposed to light for four hours and transferred in the dark growth chamber for 3 days at 22 °C. For phenotypic analysis, representative seedlings were selected for photograph. Their hypocotyl and root lengths were measured using Fiji ImageJ software.

For all imaging, RNA, ChIP, and protein related assays, *Arabidopsis* seedlings were grown on MS plates in the air-tight containers with a flow of hydrocarbon-free air at 22 °C for 3 days, and subsequently treated with ethylene gas (10 ppm) or hydrocarbon-free air for indicated hours prior to collect samples.

### Plasmid construction and generation of transgenic *Arabidopsis* plants

For yeast two-hybrid assays, the truncated *HAF2* coding regions were obtained by RT-PCR from Col-0 cDNA and then cloned into pEXPAD502 (AD) and pDBLeu (BD) backbone. For recombinant protein expression of wild type HAF2-C, HAF2M and HAF2-MC and mutated HAF2-C and HAF2-MC were cloned into pMAL-pX2 for MBP fusion. For plant transformation, full-length HAF2 cDNA was cloned into pCambia1300 to generate HAF2-FLAG-BFP and HAF2-MC-FLAG-GFP. All these genes were expressed under the control of the CaMV 35S promoter or native promoter. These constructs were introduced into *Agrobacterium tumefaciens* strain GV3101 through electroporation and transformed into *Arabidopsis* Col-0 ecotype or desired genetic backgrounds using the floral dip method. Positive T1 transformants were selected on MS medium with appropriate antibiotics. Homozygous T3 transgenic seedlings with single T-DNA insertion were used for further analysis. Supplementary Table S2 contains details of plasmid and primer sequence information.

### Yeast Two-Hybrid Assay

Yeast two-hybrid assay was performed using the ProQuest Two-Hybrid System (Invitrogen) by PEG mediated transformation following the manufacturer’s instructions. Briefly, the bait and prey plasmids were co-transformed into the yeast strain AH109 as the experiment design. The positive transformants were selected on SD/−Trp−Leu (two-dropout) medium. The growth on SD/−Trp−Leu−His (three-dropout) medium supplemented with appropriate concentrations of 3-amino-1,2,4-triazole (3-AT) indicates interaction between corresponding proteins.

### *In vitro* protein purification and pull-down assays

MBP and recombinant different version of MBP-HAF2, and 6XHis-EIN2-C proteins were expressed in the *E. coli* strain BL21-CodonPlus(DE3)-RIPL (Agilent) with protein induction by 1 mM IPTG (isopropyl b-D-1-Thiogalactopyranoside) when OD600 reached 0.5 for 4 h at 37 °C and then they were purified using amylose resin column (New England Biolabs, for MBP related proteins purification) or Ni-NTA agarose column (Qiagen, for 6XHis-EIN2-C purification) according to manufacturer’s menu followed by sonication. MBP-tagged PDC proteins or MBP proteins were added and incubated with 6XHis-EIN2-C bound Ni-NTA resin for one hour at 4 °C with gentle rocking. After five times washing with a pull-down buffer to remove non-specific binding, Ni-NTA resin beads were collected by brief centrifugation and then resuspended and denatured in 2X protein sample buffer by boiling. Proteins were then resolved by SDS–PAGE and detected with corresponding antibodies.

### Confocal imaging

Confocal images were acquired by a Zeiss LSM710 confocal microscope with a ×40 (water immersion) or ×63 len (water immersion). For other confocal imaging experiments of the co-localization of EIN2 and HAF2, 3-day-old etiolated seedlings were grown on MS with or without 10µM ACC before the images were taken from the similar middle regions of hypocotyls. The excitation/emission spectra for fluorescent proteins and fluorophores used in this study are: YFP, 514 nm/520–540 nm; BFP, 405 nm/410–500 nm.

### AlphaFold2 structure prediction

The AlphaFold2 ColabFold notebook (version 1.5.3) was used to predict the structure of the HAF2 bromodomain. The model of an acetyl-lysine residue bound to HAF2 was built using MAESTRO (version 13.5.128) from the Schrödinger suite. The acetyl-lysine was initially positioned into the active site of HAF2 by aligning the HAF2 structure to the structure of a bromodomain in complex with H3K9ac (PDB Code: 2RNW) in PyMol (version 2.4.1). The position of the acetyl-lysine was further adjusted manually into the active site of HAF2 in PyMOL. Energy minimization of the model was performed in MAESTRO by applying the OPLS_2005 force field. All structural figures were visualized using PyMol and ChimeraX (version 1.6.1).

### In vitro peptide binding assay

1 μg of purified protein, 0.1 μg of either unmodified or acetylated histone peptides, and 5 μl of Streptavidin Magnetic Beads (RayBiotech) were meticulously combined in a binding buffer (comprising 20 mM Tris-HCl pH 7.4, 200 mM NaCl, 0.6% NP-40, 1 mM EDTA, and 1 mM PMSF). The amalgamation underwent gentle rotation for 30 minutes at 4 °C. Subsequently, the protein-bound beads underwent a thorough washing procedure—three times with the binding buffer, employing gentle rotation for 5 minutes at 4 °C. The beads were then boiled in SDS-loading buffer for subsequent western blotting using MBP antibodies (NEB, E8032S). Histone peptides utilized in this study were synthesized by EpiCypher and included H3 (1–32 aa), H3K14Ac (1–20 aa), and H3K23Ac (15–34 aa).

### Histone acetyltransferase activity assay

The HAT activity assay involved the use of recombinant HAF2 variant proteins purified from E. coli as enzymes, recombinant mononucleosomes (EpiCypher) as substrates, and acetyl-CoA as a donor of acetyl groups. For the assay, 0.15 μg of enzyme and 0.5 μg of substrate were combined in HAT buffer (100 mM HEPES-NaOH pH 7.4) containing 20 μM acetyl-CoA and 50 ng/μl BSA. The reaction was conducted for 2 hours at 30 °C. The resultant reaction products were then subjected to western blotting using H3K14Ac (Millipore, 07-353) and H3K23Ac (Millipore, 07-355) antibodies.

### Co-immunoprecipitation and Immunoblot Assays

For co-immunoprecipitation (Co-IP) between EIN2-C and HAF2 following study previous briefly: Total protein extracts from plants mentioned in the paper were immunoprecipitated by DYKDDDDK Fab-Trap™ agarose beads (ChromoTek) for two hours at 4 °C with gentle rotating and then beads were washed five times with ice-cold 1XPBS buffer. The EIN2-C native antibody was prebound with the equilibrated protein-G coated magnetic beads (Dynabeads, Thermo Fisher) with 1% BSA for 3 h with gentle rotation at 4 °C and incubated at 4 °C after cleared cell fractionation addition with gentle rocking overnight for Co-IP. Dynabeads was magnetically precipitated using a DynaMagnetic rack (Thermo Fisher) and then washed five times with 0.5 mL of ice-cold 1XPBS buffer to eliminate non-specific binding. After IP and washing, proteins were then released from beads using 2× Laemmli sample buffer by heating at 85 °C for 8 min and subject to SDS-PAGE. For EIN2 related western blots, protein samples were denatured in 2× Laemmli sample buffer with the addition of 3M urea at 75 °C in water bath for 5 min to avoid protein aggregation and degradation.

For protein immunoblots, proteins were separated by SDS–PAGE and transferred to a nitrocellulose membrane (0.2 µm, Bio-rad) by the wet-tank transfer method and blocked with 5% non-fat milk in TBST for 1 h at room temperature prior to the overnight inoculation in primary antibody at 4 °C with gentle rotation. The following antibodies and dilutions were used for immunoblotting: anti-HA (Biolegend, #901503), 1:10000; anti-FLAG (rabbit) (CST, #14793), 1:2000; and anti-FLAG (mouse) (Sigma, F3165),1:5000; anti-H3 (abcam, ab1791), anti-EIN2-C (developed in-house)^18^, 1:2000. Goat anti-mouse Kappa Light Chain antibody (Bio-Rad, #105001G) and Goat Anti-Rabbit (IgG (H + L)-HRP Conjugate #1706515, Bio-Rad) were used as secondary antibodies at 1:10000 dilution. HRP activity was detected by enhanced chemiluminescence (ECL, GE Healthcare) according to the manufacturer’s instructions using either a ChemiDoc Imaging System (Bio-Rad) or conventional X-ray films.

### CRISPR-Cas9 mutagenesis

In brief, pairs of two guide RNA (gRNA) sequences targeting *HAF2* locus were designed by using CRISPR-PLANT v2 (http://omap.org/crispr2/index.html) online tool. gRNA sequences with high specificity in *Arabidopsis* genome and high predicted efficiency were chosen for PCR amplification using pDT1T2 vector as template and later incorporated into the binary vector pHEE401E. *Agrobacterium* mediated floral dipping method was used to transform the *HAF2* gRNA containing pHEE401E vectors into each corresponding genetic background. To genotype deletion mutations, plant genomic DNAs extracted from hygromycin positive T1 plants were used to PCR amplify of *HAF2* gRNA targeted regions and followed by Sanger sequencing to identify the CRISPR-Cas9 directed mutation and to obtain homozygotes of *haf2* mutations in Col-0 or *e1-2-2e2-2*.

### Transcriptome analysis

RNA extraction was conducted according to the manufacturer’s recommendation using the RNeasy Plant Kit (Qiagen) with DNase I digestion and clean up (Qiagen, RNeasy Mini Kit (250)). For mRNA-seq, libraries were generated according to the instructions from manufacturer using the NEBNext® Poly(A) mRNA Magnetic Isolation Module (E7490) and the NEBNext® Ultra™ II RNA Library Prep Kit for Illumina® (E7770) with 1 μg RNA as input. Indexed libraries were sequenced on the platform of HiSeq2000 (Illumina). Paired-end raw fastq data were evaluated with FastQC and low-quality reads were removed with Trim Galore (Babraham Institute). The trimmed and filtered reads were then mapped to the *Arabidopsis* reference genome (TAIR10) with STAR ^74^. Mapped reads were counted by featureCounts (Subread 2.0.1) and differentially regulated genes were identified using R package DESeq2 ^75^ with a p-adj< 0.05 and | log2(fold change) | > 1. The RNA-seq data for Col-0 air and ethylene treatment were obtained from previous study (GEO accession code GSE77396) ^24^ and reanalyzed with the pipeline described above. The scatterplots, violin plots, and dot plots were generated by R package *ggplot2* and the heatmaps of log2FC were generated by R package *pheatmap*. GO enrichment analysis for DEGs was performed by agriGO v2 ^76^.

### Histone extraction

Approximately two grams of ground fine powders of etiolated seedlings were homogenized in 20 mL nuclear extraction buffer (250 mM sucrose, 60 mM KCl, 15 mM NaCl, 5 mM MgCl_2_, 1 mM CaCl_2_, 15 mM PIPES pH = 6.8, 0.5% Triton X-100, 1 mM PMSF, 1X protease inhibitor, and 10 mM sodium butyrate as HDAC inhibitor) for 10min. After filtering through a single layer of Miracloth twice, the flow-through was then centrifuged at 2880Xg for 20 min at 4°C to collect the nuclei. The crude nuclear pellet was then resuspended in 800 μL 0.4 M H_2_SO_4_ by gentle pipetting, and was incubated for two hours for complete lysis followed by centrifuge at 16000Xg at 4 °C. Total histone in the supernatant phase was then precipitated by 33% trichloroacetic acid (TCA) for at least two hours and then centrifuged at maximum speed for 10 min at 4 °C. The histone pellet was washed twice with ice-cold acetone containing 0.1% HCl and once with pure acetone. The air-dried histone pellet was then dissolved in 2X Laemmli buffer containing 2 M urea, denatured by boiling, and subject to immunoblots.

### ChIP-seq analysis

Chromatin immunoprecipitation was performed according to previous publication ^22^. In brief, 3-day-old etiolated *e1-2-2 e2-2 #17* seedlings treated with air or 4h ethylene were harvested and crosslinked in 1% formaldehyde, and the chromatin was isolated and sheared by Bioruptor. Anti-H3K14ac (Millipore; 07-353) and anti-H3K23ac (Millipore; 07-355) antibodies together with Dynabeads were added to the sonicated chromatin followed by incubation overnight to precipitate histone bound DNA fragments. DNA eluted from Chromatin-immunoprecipitated was prepared for library construction by using NEBNext® Ultra™ II DNA Library Prep with Sample Purification Beads according to standard operating procedures and then sequenced using HiSeq2000 (Illumina). Paired-end raw reads were first mapped to the *Arabidopsis* genome (TAIR10) using Bowtie2 software with default parameters. Duplicate reads were removed using SAMtools. H3K14ac and H3K23ac ChIP-seq data for Col-0 air and ethylene treatment were obtained from previous study (GEO accession code GSE77396) ^24^. ChIP-seq coverage tracks shown in the genome browser IGV were generated using deepTools 3.5.0 ^77^ bamCoverage function normalized by reads per genomic content (1x normalization, 1X RPGC) and bin size = 1. The ChIP-seq signals from 2kb upstream TSS to 2kb downstream TES of PDC specifically regulated gene were calculated with bamCompare and computeMatrix (deepTools), and plotted with plotHeatmap and plotProfile in deepTools or *ggplot2* in R.

## Acknowledgements

We thank D. Hernandez for lab maintenance. We thank P. Oliphint, A. Webb, and Microscopy and Imaging Facility at The University of Texas at Austin for the support of confocal microscope imaging. We thank *Arabidopsis* Biological Resource Center for the *Arabidopsis* T-DNA insertion mutants. This work was supported by grants from the National Institute of Health to H.Q. (NIH-2R01 GM115879) and Lorene Morrow Kelley Endowed Faculty Fellowship Fund from UT Austin.

## Author contributions

Chia-Yang Chen and H.Q. designed the study. Chia-Yang Chen, Z.S. performed most of the experiments and analysis. Bo Zhao cloned HAF2 cDNA, G. W performed ChIP-seq and RNA-seq data analysis. J Z. and H.H performed structure prediction, J.G.B. and P.K. helped genetic material generation and imaging. H.Q. supervised the study. H.Q. and Chia-Yang Chen wrote the paper. **Competing interests**: The authors declare no competing interests. **Data and materials availability**: Further information and requests for all unique materials generated in this study may be directed to and will be fulfilled by the corresponding author Hong Qiao. The high-throughput sequencing data generated in this study have been deposited in the Gene Expression Omnibus (GEO) database (GSE249043).

